# Tobramycin adaptation alters the antibiotic susceptibility of *Pseudomonas aeruginosa* quorum sensing-null mutants

**DOI:** 10.1101/2023.01.13.523864

**Authors:** Rhea G. Abisado-Duque, Kade A. Townsend, Brielle M. Mckee, Kathryn Woods, Pratik Koirala, Alexandra J. Holder, Vaughn D. Craddock, Matthew Cabeen, Josephine R. Chandler

**Author notes:** Contributed equally (Rhea G. Abisado-Duque, Kade A. Townsend, Brielle M. Mckee). Address correspondence to Josephine R. Chandler,.

## Abstract

The opportunistic bacterium *Pseudomonas aeruginosa* uses the LasR-I quorum sensing system to increase resistance to the aminoglycoside antibiotic tobramycin. Paradoxically, *lasR*-null mutants are commonly isolated from chronic human infections treated with tobramycin, suggesting there may be a mechanism allowing the *lasR*-null mutants to persist under tobramycin selection. We hypothesized that the effects of inactivating *lasR* on tobramycin resistance might be dependent on the presence or absence of other gene mutations in that strain, a phenomenon known as epistasis. To test this hypothesis, we inactivated *lasR* in several highly tobramycin-resistant isolates from long-term evolution experiments. We show that the effects of Δ*lasR* on tobramycin resistance are strain dependent, which is due to a single mutation in the *fusA1* gene encoding the translation elongation factor EF-G1A (G61A nucleotide substitution). The *fusA1* G61A mutation confers a strong selective advantage to Δ*lasR* mutants under tobramycin treatment. The effects of *fusA1* G61A on Δ*lasR-*dependent tobramycin resistance are dependent on the MexXY efflux pump and the MexXY regulator ArmZ. The *fusA1* mutation also modulates Δ*lasR* mutant resistance to two other antibiotics, ciprofloxacin and ceftazidime. Our results provide a possible explanation for the emergence of *lasR-*null mutants in clinical isolates and illustrate the importance of epistatic gene interactions in the evolution of quorum sensing.

## IMPORTANCE

*Pseudomonas aeruginosa* causes chronic infections in the airways of patients with the genetic disease cystic fibrosis. In these clinical isolates the quorum-sensing regulator LasR frequently becomes mutated. The emergence of LasR-null mutants can have important consequences to clinical outcomes. We performed studies of tobramycin-resistant isolates from laboratory evolution experiments to understand the factors that contribute to the emergence of LasR mutants. Although mutating *lasR* was deleterious in our laboratory strain, we observed that this same mutation was beneficial in some of the isolates we tested. This difference was due to a single mutation in the *fusA1* gene encoding the translation elongation factor EF-G1A (nucleotide G61A). We show that *fusA1* G61A confers an advantage to *lasR* mutants under tobramycin selection. Thus in populations undergoing adaptation to new environments, some mutations could arise that permit the emergence of *lasR-*null mutations and ultimately alter the course of infection.

## INTRODUCTION

*Pseudomonas aeruginosa* is an opportunistic multidrug resistant pathogen that regulates about 10% of its genes using quorum sensing, a type of bacterial communication that is activated in a population density-dependent manner (1-4). One type of quorum sensing involves a LuxR-family signal receptor and a LuxI-family signal synthase (2, 5, 6). In *P. aeruginosa,* there are two complete LuxI-R-type quorum-sensing systems; the LasI-R system, which produces and responds to the signal *N*-3-oxododecanoyl-homoserine lactone (3OC12-HSL) (2-4,7) and the RhlI-R system, which produces and responds to the signal *N*-butanoyl-L-homoserine lactone (C4-HSL) (8). In laboratory strains, the LasR-I system regulates expression of the *rhlR* and *rhlI* genes, resulting in a hierarchy in quorum sensing, wherein LasR is the master regulator (5, 7, 9). The LasI-R system is essential for pathogenesis in several acute infection animal models (10-12).

We, and others, have shown that the LasR-I system increases *P. aeruginosa* resistance to tobramycin, a clinically relevant antibiotic that is commonly used to treat acute and chronic *P. aeruginosa* infections (13-17). Quorum-sensing systems in other bacteria can also increase antibiotic resistance (18), suggesting that the control of antibiotic resistance by quorum sensing might provide some advantages that are conserved across several bacterial species and environments. The *P. aeruginosa* LasR-I system increases tobramycin resistance in planktonic cultures (17) and also in biofilm conditions (13-16). Tobramycin can also suppress the proliferation of *lasR* mutants that emerge de novo during propagation of populations in certain conditions in the laboratory (17). These results suggest tobramycin treatment could have a strong selective pressure on quorum sensing in a clinical setting.

Clinical isolates of *P. aeruginosa* are very genetically diverse in part due to adaptations to the cystic fibrosis lung environment (19-21). One of the most commonly observed adaptations in clinical infections is *lasR* mutation, resulting in the loss of quorum-sensing function (21-25). Although LasR contributes to pathogenesis in acute infection models with laboratory strains (26-28), the emergence of LasR mutants in a clinical setting correlates with worse outcomes in infection (22, 29-32). It is counterintuitive that *lasR*-null mutations emerge in infections of patients treated with tobramycin when these mutations increase sensitivity to tobramycin in laboratory strains. There are several potential explanations for this puzzling finding. Certain nutritional conditions can select for *lasR* mutants (23, 33-35), which could possibly be stronger selection than tobramycin. Alternatively, certain conditions might alter the physiology of *lasR* mutants to make them more tobramycin resistant (36, 37). Whether the conditions of the infection environment contribute to the selection of *lasR* mutations in infections of tobramycin-treated patients remains unknown.

Here, we sought to explore another possible explanation for *lasR* mutant emergence in infections of tobramycin-treated patients; that the effects of tobramycin on *lasR*-null mutants might be dependent on genetic background. In particular, we hypothesized that tobramycin susceptibility of Δ*lasR* mutants might be modified by epistatic gene interactions (38). We identify one such mutation; a point mutation in the gene encoding the translation accessory factor EF-G1A (*fusA1*). We show that the *fusA1* G61A (FusA1^A21T^) mutation facilitates the emergence of Δ*lasR* mutations in populations under tobramycin selection. Our results show that antibiotic susceptibility of *lasR* mutants can be genotype dependent and support the idea that epistatic interactions could contribute to the emergence of LasR-null mutations in a clinical setting.

## RESULTS

### Δ*lasR* has strain-dependent effects on tobramycin resistance

We previously characterized six tobramycin-resistant genetic isolates of *P. aeruginosa* PA14 with mutations in unique genes (e.g. *fusA1* and *ptsP*, see Table S1)(17). We used these isolates to test the hypothesis that antibiotic adaptations could alter the effect of Δ*lasR* mutations on antibiotic resistance. To test this hypothesis, we used allelic exchange to delete *lasR* from each of the six isolates from our prior study (termed T1-T6). We compared the minimum inhibitory concentration (MIC) of the Δ*lasR* isolates with their parent (Fig. 1A). Similar to our prior results (17), deleting *lasR* caused a small decrease in tobramycin resistance of PA14, although in this study the difference was not significant due to comparisons with a wider range of MICs in our statistical analyses. Deleting *lasR* in the T1 isolate increased tobramycin resistance ∼2-fold (Fig. 1A). Deleting *lasR* in T3 had a smaller, but similar effect (Fig. 1A). In pairwise comparisons with the PA14 wild-type strain, the effects of Δ*lasR* on tobramycin MIC were significantly different in T1 and T3 (p<0.05, Fig. 1A). We did not observe significantly different effects of deleting *lasR* on the MIC for any of the other 4 isolates (Fig. S1).

**Fig. 1.**
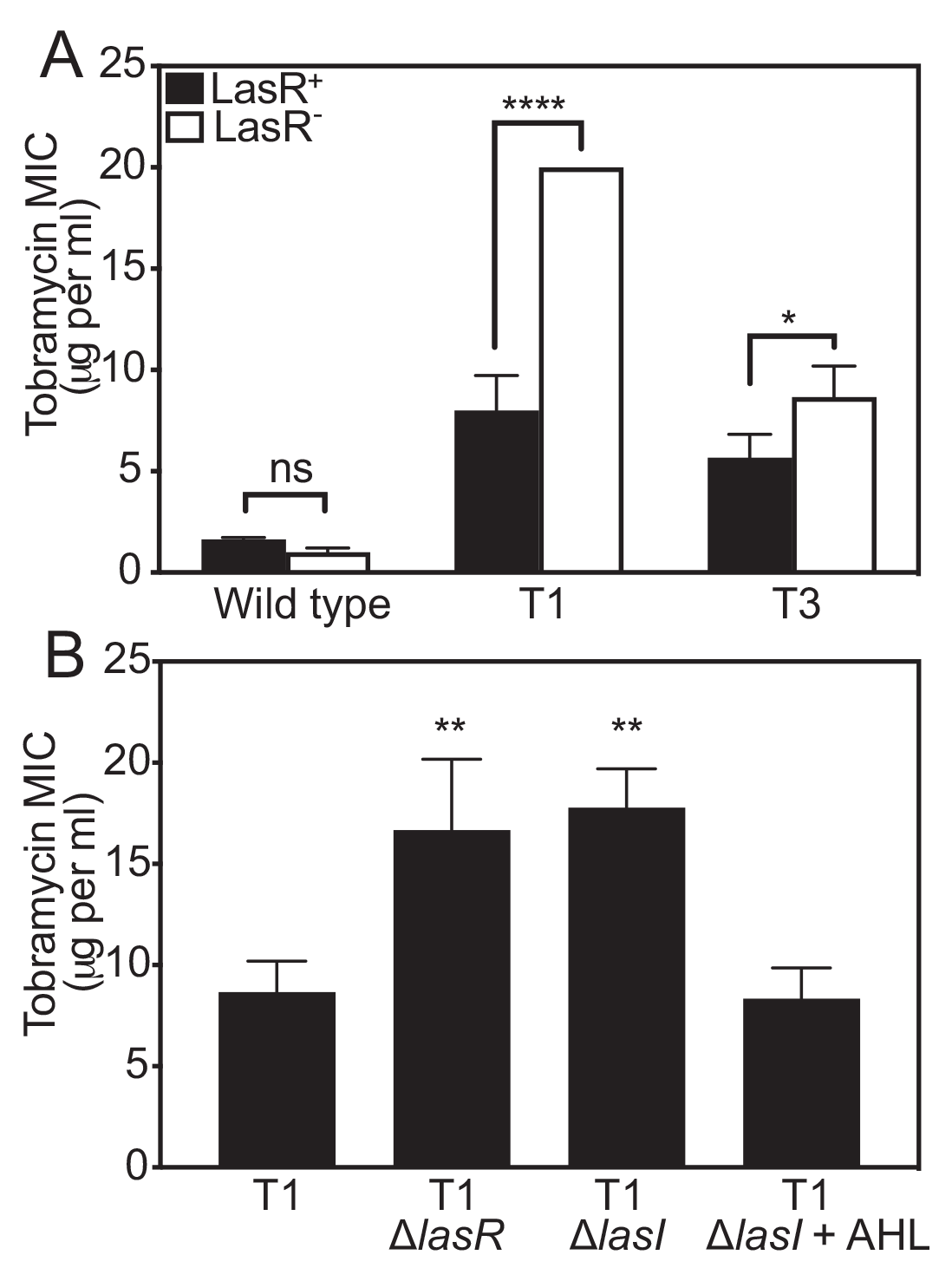
Tobramycin resistance of Δ*lasR* mutants is strain dependent. **A.** The minimum inhibitory concentration (MIC) of tobramycin was determined for each strain carrying intact *lasR* (LasR^+^, black bars) or Δ*lasR* (LasR^-^, white bars). The significance of the effect of strain (wild type, T1 or T3) and *lasR* allele (LasR^+^ or LasR^-^) and the interaction of the two on MIC was determined by 2-way ANOVA by performing pair-wise comparisons of each LasR^+/-^ strain pair with that of PA14. The interaction was significant for T1 (F_1,8_ = 156.8 and p<0.0001) and T3 (F_1,8_ = 10.67 and p < 0.05). Pairwise comparisons of LasR^+^ and LasR^-^ of each strain were determined using Sidak’s post-hoc test with p-values adjusted for multiple comparisons; *, p<0.05; ****, p < 0.0001; ns = not significant. **B.** Tobramycin MIC of T1 and T1-mutated strains. Where indicated, AHL (3OC12-HSL) was added prior to inoculating culture tubes at 10 µM final concentration. Statistical analysis was by one-way ANOVA and Dunett’s multiple comparisons with T1; **, p<0.01. For both panels, results are the average of three independent experiments and the vertical bars indicate standard deviation.

To further confirm the effects of the LasR-I system on tobramycin resistance in the T1 isolate, we deleted the *lasI* signal synthase gene (Fig. 1B). Because LasI synthesizes the LasR signal, we anticipated that deleting *lasI* would cause changes in tobramycin MIC similar to that of a *lasR* deletion. Consistent with this expectation, deleting *lasI* significantly increased tobramycin resistance in T1, and adding synthetic 3OC12-HSL to the Δ*lasI* mutant culture restored resistance levels to that of T1 (Fig. 1B). These results offer further support that the effects of disrupting the LasR-I circuit on tobramycin resistance are strain dependent.

### *fusA1* (G61A) modulates the effects of Δ*lasR* on tobramycin resistance

We hypothesized that one or more genetic mutations in T1 and T3 altered the susceptibility of Δ*lasR* mutants to tobramycin. We previously described that the T1 and T3 isolates have only two common mutations, *fusA1* G61A (FusA1^A21T^) and *ptsP* 1547T (a frameshift mutation that is predicted to result in an inactivated protein) (Table S1)(17). We individually introduced each of these mutations to PA14 and PA14 Δ*lasR* using allelic exchange and compared the MIC of the mutated strains to that of the PA14 parent strains (Fig. 2A). We found that deleting *lasR* in PA14 *fusA1* G61A increased the MIC of the *fusA1* G61A mutant ∼2- fold, which was a significantly different effect than deleting *lasR* in PA14 (p < 0.0001). In contrast, deleting *lasR* in the *ptsP* 1547T mutant was not significantly different from that of PA14 (p=0.9865). These results suggested that the *fusA1* G61A mutation somehow increases tobramycin resistance of Δ*lasR* mutants. We next asked if ectopic expression of the wildtype *fusA1* would restore the MIC to that of wild type. To do so, we fused the wild-type *fusA1* gene to a rhamnose-inducible promoter (P*rha*) and introduced this cassette to the neutral *attB* site in the genome of T1 and the PA14 *fusA1* G61A mutant (Fig. 2B and Fig. S2). In wild-type *fusA1* expressing strains, the effects of deleting *lasR* on the tobramycin MIC were indistinguishable from that of the parent (T1 or PA14 *fusA1* G61A carrying P*rha-fusA1* compared with Δ*lasR* of each strain). Thus, the effects of the *fusA1* G61A mutation on *lasR-*dependent tobramycin resistance can be restored by complementation with wild type *fusA1*.

**Fig. 2.**
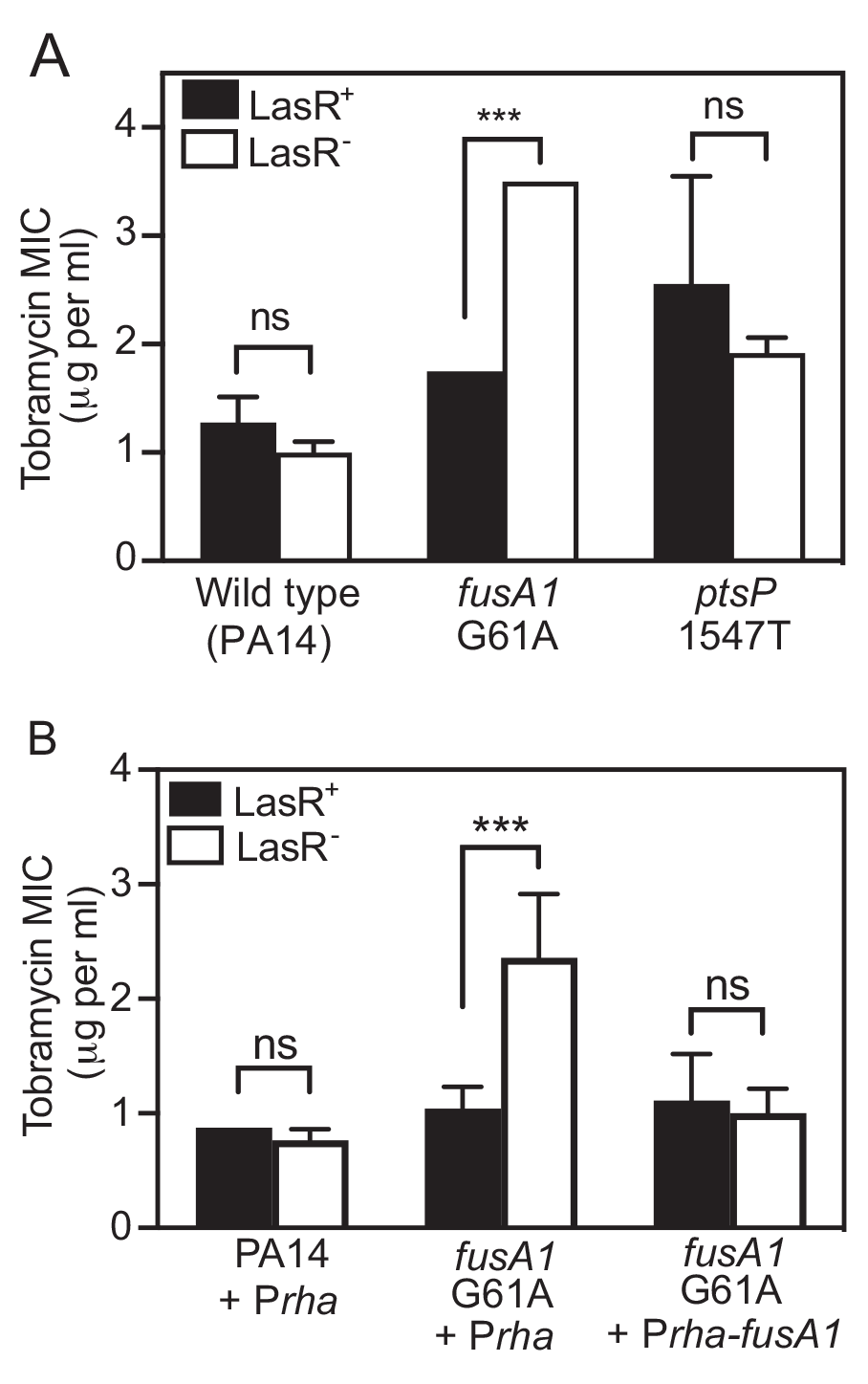
Effects of *fusA1* G61A and Δ*lasR* on tobramycin resistance. The minimum inhibitory concentration (MIC) of tobramycin was determined for each strain carrying intact *lasR* (LasR^+^, black bars) or Δ*lasR* (LasR^-^, white bars). The significance of the effect of strain (indicated along the X axis) and *lasR* allele (LasR^+^ or LasR^-^) and the interaction of the two on MIC was determined by 2-way ANOVA by performing pair-wise comparisons of each LasR^+/-^ strain pair with that of PA14 (A) or PA14 P*rha* (B). **A.** There was a statistically significant interaction between the effects of strain and *lasR* for *fusA1* G61A (F_1,8_ = 156.8, p < 0.0001) but not for *ptsP* 1547T (F_1,8_ = 0.3642, p = 0.5629). **B.** Strains had either the empty CTX-1 P*rha* or the CTX-1 P*rha-fusA1* cassette inserted into the neutral *attB* site in the genome. Rhamnose was added to all cultures at 0.4% final concentration. There was a statistically significant interaction between the effects of strain and *lasR* for *fusA1* G61A P*rha* (F_1,8_ = 17.41, p < 0.005) but not for *fusA1* G61A + P*rha-fusA1* (F_1,8_ = 1.7×10^-7^, p = 0.997). For both panels, results are the average of three independent experiments and the vertical bars indicate standard deviation. Pair-wise comparisons of LasR^+^ and LasR^-^ of each strain were determined using Sidak’s post-hoc test with p-values adjusted for multiple comparisons; ***, p<0.001; ns = not significant.

### Tobramycin has strain-dependent effects on the proliferation of Δ*lasR* mutants in cocultures

We sought to assess the selective effects of tobramycin on Δ*lasR* mutants in different strain backgrounds using coculture competition experiments. When PA14 is co-inoculated with PA14 Δ*lasR*, the Δ*lasR* mutant rapidly proliferates to higher frequency because it has a fitness advantage over PA14; however, Δ*lasR* mutants are suppressed in identical cocultures grown with subinhibitory tobramycin (17). We hypothesized that tobramycin would enhance rather than suppress the proliferation of Δ*lasR* mutants in strains carrying the *fusA1* G61A mutation. To test this hypothesis, we inoculated cocultures with either PA14 or PA14 *fusA1* G61A and a 1% initial population of each respective Δ*lasR* mutant and grew the cocultures with or without tobramycin. Cocultures were transferred daily to fresh medium and the proportion of Δ*lasR* in the final population was assessed after 2 daily transfers. The results (Fig. 3) showed that tobramycin had significantly different effects on Δ*lasR* mutant proliferation in each of the two strain backgrounds (p<0.0001). At the end of the PA14 experiment the Δ*lasR* mutants were ∼30% of the population in cocultures grown with no tobramycin but reached only ∼0.5% of the tobramycin-treated population (Fig. 3, left), consistent with prior results (17). In cocultures with strains carrying *fusA1* G61A, the Δ*lasR* mutants similarly reached ∼30% of the population in the absence of tobramycin but further increased to ∼60% of the population in the presence of tobramycin (Fig. 3, right). These results support the idea that under tobramycin selection, the *fusA1* G61A mutation causes Δ*lasR* mutations to become beneficial, rather than deleterious, to bacterial fitness.

**Fig. 3.**
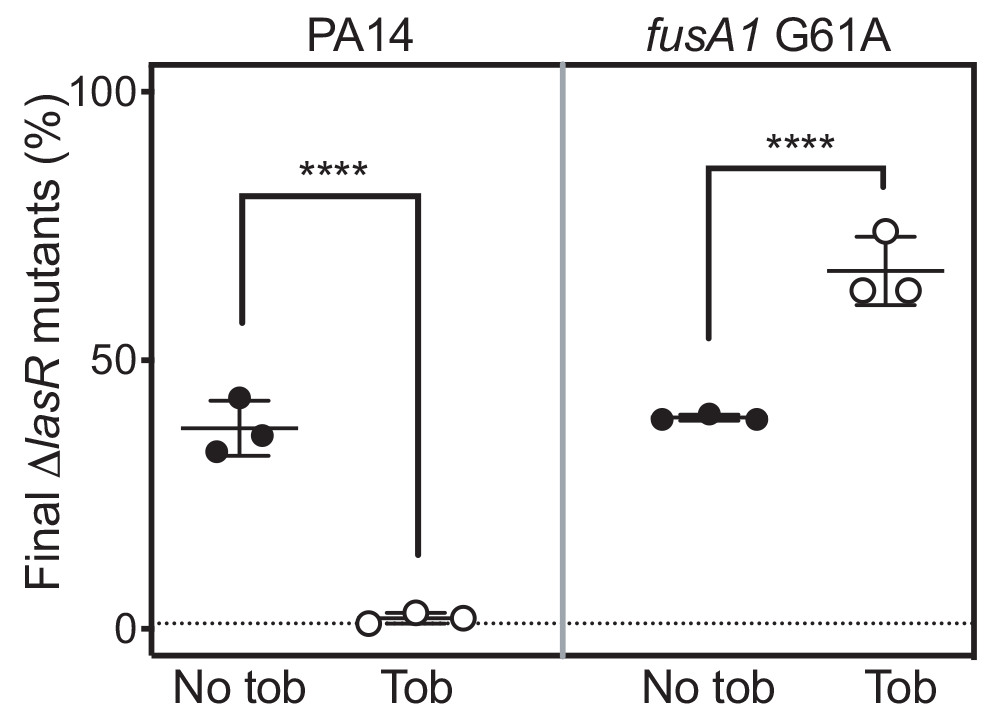
Tobramycin effects on Δ*lasR* proliferation in cocultures. Cocultures of wild-type PA14 and PA14 Δ*lasR* (left panel), or of *fusA1* G61A and *fusA1* G61A Δ*lasR* (right panel) were grown with no tobramycin (No tob, closed circles) or with tobramycin (Tob, open circles) at the highest concentration that permits growth (0.3 μg/ml for PA14 or 2 μg/ml for *fusA1* G61A). In both cases the Δ*lasR* mutant was started at 1% of the total coculture population. After combining, cocultures were inoculated into casein medium and subsequently transferred to fresh medium daily for two days. Final population densities ranged from 1-5 x 10^9^ cells per ml. Each data point represents a single experiment and vertical lines represent standard deviation. Statistical analysis by 2-way ANOVA was used to determine the significance of the effect of the tobramycin and the *lasR* allele and interaction between the two on the relative fitness of the Δ*lasR* mutant and the interaction was significant (p < 0.0001 and F_1,8_ = 41.26). Pair-wise comparisons of LasR^+^ and LasR^-^ for each condition (+/- tob) were performed using a Sidak’s post-hoc analysis with p-values adjusted for multiple comparisons; ****, p < 0.0001.

### Effects of other *fusA1* mutations on tobramycin MIC of Δ*lasR* mutants

The *fusA1* gene encodes elongation factor G1A (EF-G1A), a GTPase protein that hydrolyzes GTP to drive the elongation and recycling steps of protein synthesis (39, 40). EF-G1A is an essential component in translation, and *fusA1*-null mutations are not viable (39-41). Nevertheless, *fusA1* has been reported to be among the most frequently mutated genes in clinical isolates (42-45), and in at least some cases *fusA1* mutations increase tobramycin resistance (41, 42, 46). EF-G1A has 5 domains (labeled I-V in Fig. 4A). The FusA1 A21T substitution is within a motif in domain 1 called the Walker-A P-loop, which is responsible for binding phosphoryl groups and catalyzing phosphoryl transfer of NTPs (47). FusA1^A21T^ has not been reported in the literature to our knowledge, however, a survey of 4,312 sequenced strains identified 4 strains with this mutation (Table S1), indicating *fusA1* G61A mutations do occur naturally but are relatively rare.

**Fig. 4.**
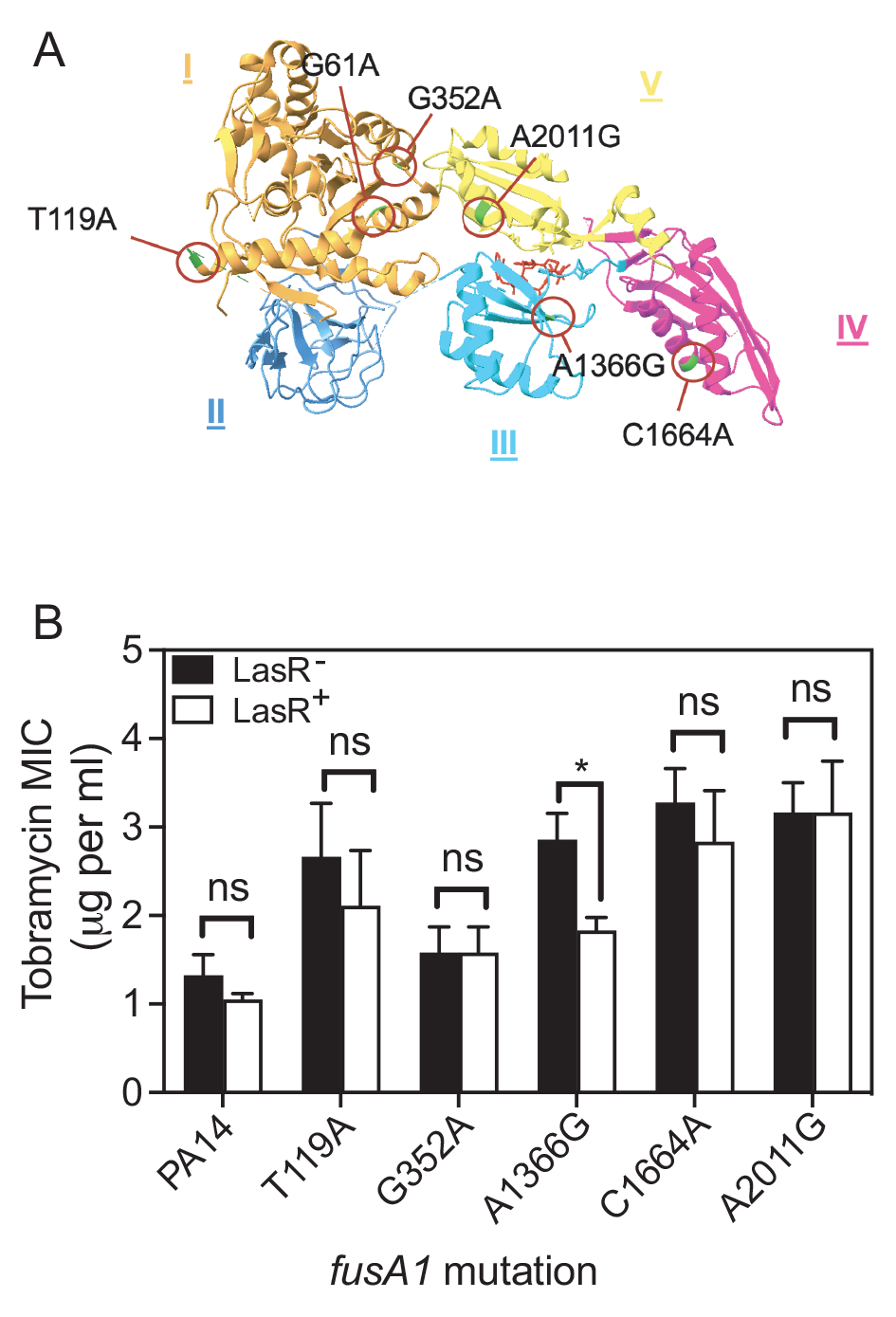
Other *fusA1* mutations and their effects on tobramycin resistance of Δ*lasR* mutants. **A.** Ribbon diagram of the *fusA1-*encoded protein Elongation Factor-G (EF-G) (PDB ID: 4FN5). The protein was crystallized bound to argyrin, which is shown in red. Each of the mutations assessed in panel B are indicated with green coloration in the ribbon diagram and a red circle and were identified using the UCSF ChimeraX (76) **B.** The minimum inhibitory concentration (MIC) of tobramycin was determined for each strain carrying the wild type *fusA1* allele (PA14) or PA14 with the indicated *fusA1* nucleotide mutation. Each strain had either an intact *lasR* (LasR^+^, black bars) or Δ*lasR* (LasR^-^, white bars). Results are the average of three independent experiments and the vertical bars indicate standard deviation. The significance of the effect of the *fusA1* allele and the *lasR* allele and the interaction of the two on MIC was determined by 2-way ANOVA by performing pair-wise comparisons of each LasR^+/-^ strain pair with that of PA14. Only A1366G showed a significant interaction (p<0.05, F_1,8_ = 10.25). Pairwise comparisons of LasR^+^ and LasR^-^ of each strain was determined using Sidak’s post-hoc test with p-values adjusted for multiple comparisons. *, p<0.05.

Several other mutations in *fusA1* have been characterized (Fig. 4A)(41). To test whether these mutations enhance or alter tobramycin resistance of Δ*lasR* mutants, we introduced five *fusA1* mutations to the genome of PA14 or PA14 Δ*lasR* by allelic exchange and determined the tobramycin MIC of each strain (Fig. 4B). In all but one (A1366G) of the *fusA1-*mutated strains, the effect of LasR on antibiotic resistance was not significantly altered compared with that of PA14. In *fusA1* A1366G, the Δ*lasR* mutation decreased tobramycin resistance compared with the LasR^+^ strain, which is the opposite of the effect observed with the G61A mutation. These results show that not all *fusA1* mutations modulate tobramycin resistance of Δ*lasR* mutants and that different mutations have independent effects.

### *fusA1* G61A modulates Δ*lasR* resistance to other antibiotics

We next asked whether the *fusA1* G61A mutation can also alter the susceptibility of Δ*lasR* mutants to antibiotics other than tobramycin. We tested the antibiotics ciprofloxacin (DNA gyrase inhibitor), ceftazidime (cell wall biosynthesis inhibitor) piperacillin (cell wall biosynthesis inhibitor), and tetracycline (ribosome inhibitor) against PA14 and our *fusA1* and *lasR* single and double mutant strains (Table 1). We observed significant strain-dependent effects of the Δ*lasR* mutation on the MIC values for two of these antibiotics: ciprofloxacin (p < 0.01) and ceftazidime (p < 0.0001). With ciprofloxacin, Δ*lasR* increased resistance of the *fusA1* G61A mutant but this mutation caused no changes in the PA14 MIC. With ceftazidime we observed different effects: the Δ*lasR* mutation decreased resistance in PA14 but not in the *fusA1* G61A mutant. These results show that *fusA1* G61A mutations modulate susceptibility of Δ*lasR* mutants to several different classes of antibiotics.

**Table 1.**
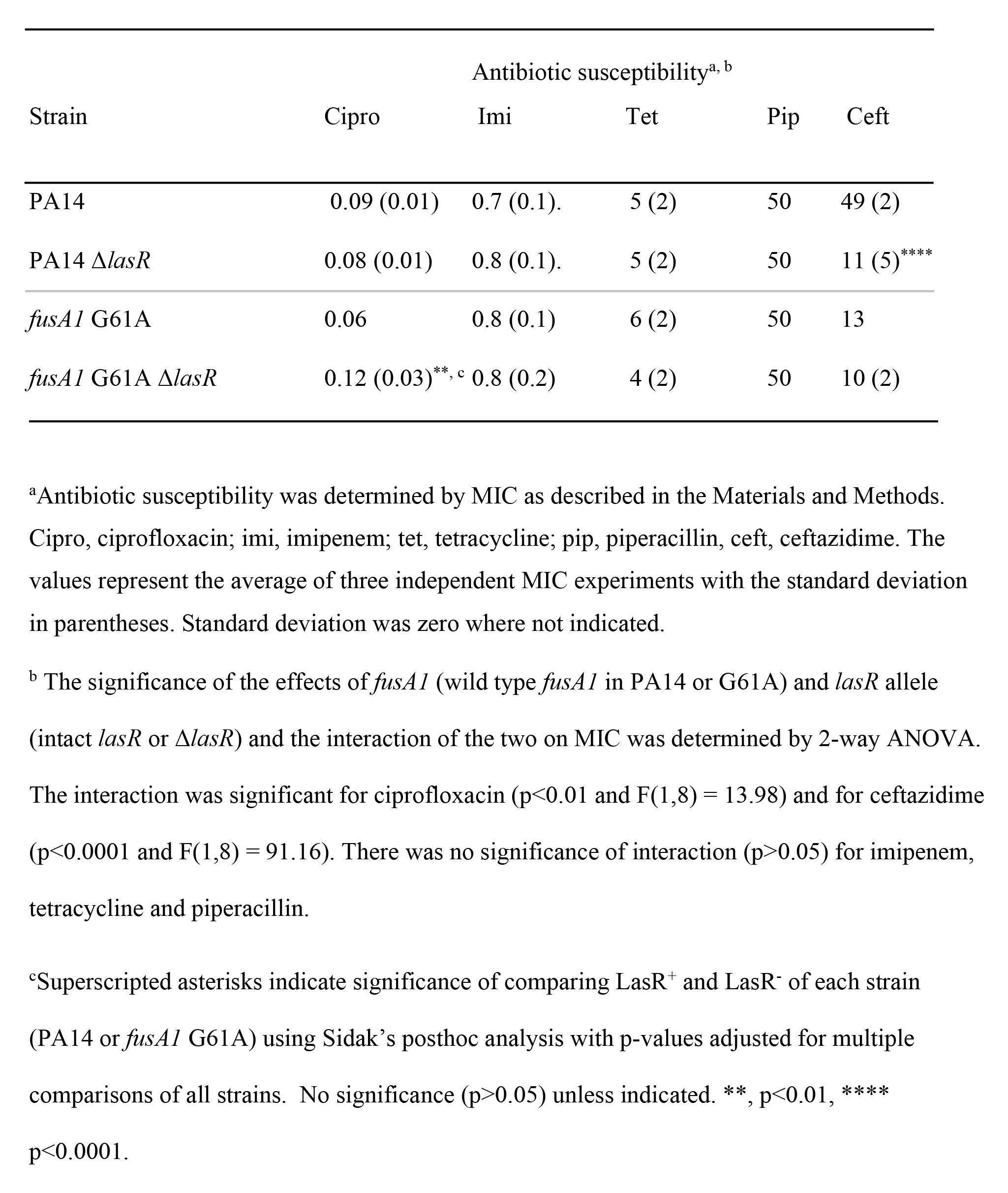
Antibiotic susceptibility of *lasR* and *fusA1* G61A mutant strains.

| Strain | Antibiotic susceptibility<sup>a, b</sup> |  |  |  |  |
| --- | --- | --- | --- | --- | --- |
|  | Cipro | Imi | Tet | Pip | Ceft |
| PA14 | 0.09 (0.01) | 0.7 (0.1). | 5 (2) | 50 | 49 (2) |
| PA14 $\Delta lasR$ | 0.08 (0.01) | 0.8 (0.1). | 5 (2) | 50 | 11 (5)<sup>****</sup> |
| *fusAI* G61A | 0.06 | 0.8 (0.1) | 6 (2) | 50 | 13 |
| *fusAI* G61A $\Delta lasR$ | 0.12 (0.03)<sup>**, c</sup> | 0.8 (0.2) | 4 (2) | 50 | 10 (2) |

<sup>a</sup>Antibiotic susceptibility was determined by MIC as described in the Materials and Methods. Cipro, ciprofloxacin; imi, imipenem; tet, tetracycline; pip, piperacillin, ceft, ceftazidime. The values represent the average of three independent MIC experiments with the standard deviation in parentheses. Standard deviation was zero where not indicated.
<sup>b</sup> The significance of the effects of *fusAI* (wild type *fusAI* in PA14 or G61A) and *lasR* allele (intact *lasR* or $\Delta lasR$ ) and the interaction of the two on MIC was determined by 2-way ANOVA. The interaction was significant for ciprofloxacin ( $p < 0.01$ and $F(1,8) = 13.98$ ) and for ceftazidime ( $p < 0.0001$ and $F(1,8) = 91.16$ ). There was no significance of interaction ( $p > 0.05$ ) for imipenem, tetracycline and piperacillin.
<sup>c</sup>Superscripted asterisks indicate significance of comparing LasR<sup>+</sup> and LasR<sup>-</sup> of each strain (PA14 or *fusAI* G61A) using Sidak's posthoc analysis with p-values adjusted for multiple comparisons of all strains. No significance ( $p > 0.05$ ) unless indicated. \*\*, $p < 0.01$ , \*\*\*\* $p < 0.0001$ .

### *fusA1* G61A and Δ*lasR* increase tobramycin resistance by activating the MexXY efflux pump

Some other *P. aeruginosa fusA1* mutations can enhance aminoglycoside resistance through the multidrug efflux pump MexXY (41, 48). Thus, we hypothesized that MexXY is important for increasing the tobramycin resistance of *fusA1* G61A and Δ*lasR* double mutants. As an initial test of this hypothesis, we measured expression of the *mexX* gene, which is immediately upstream of and in the same operon as *mexY* (Fig. 5A). We quantified *mexX* transcripts in logarithmically growing cells of PA14 *fusA1* G61A and Δ*lasR* single and double mutants. Consistent with our hypothesis, *mexX* transcripts were the highest in *fusA1* G61A and Δ*lasR* double mutants. We also deleted the *mexY* gene that is essential for pump activity. Our attempts to delete *mexY* in *fusA1* G61A were unsuccessful; however, we were able to delete this gene in T1 and T1 Δ*lasR*, and also PA14 and PA14 Δ*lasR*. Deleting Δ*mexY* in T1 abolished the LasR-dependent changes in MIC observed in this strain (Fig. 5B), supporting that MexY is important for *fusA1* G61A to increase tobramycin resistance of Δ*lasR* mutants.

**Fig. 5.**
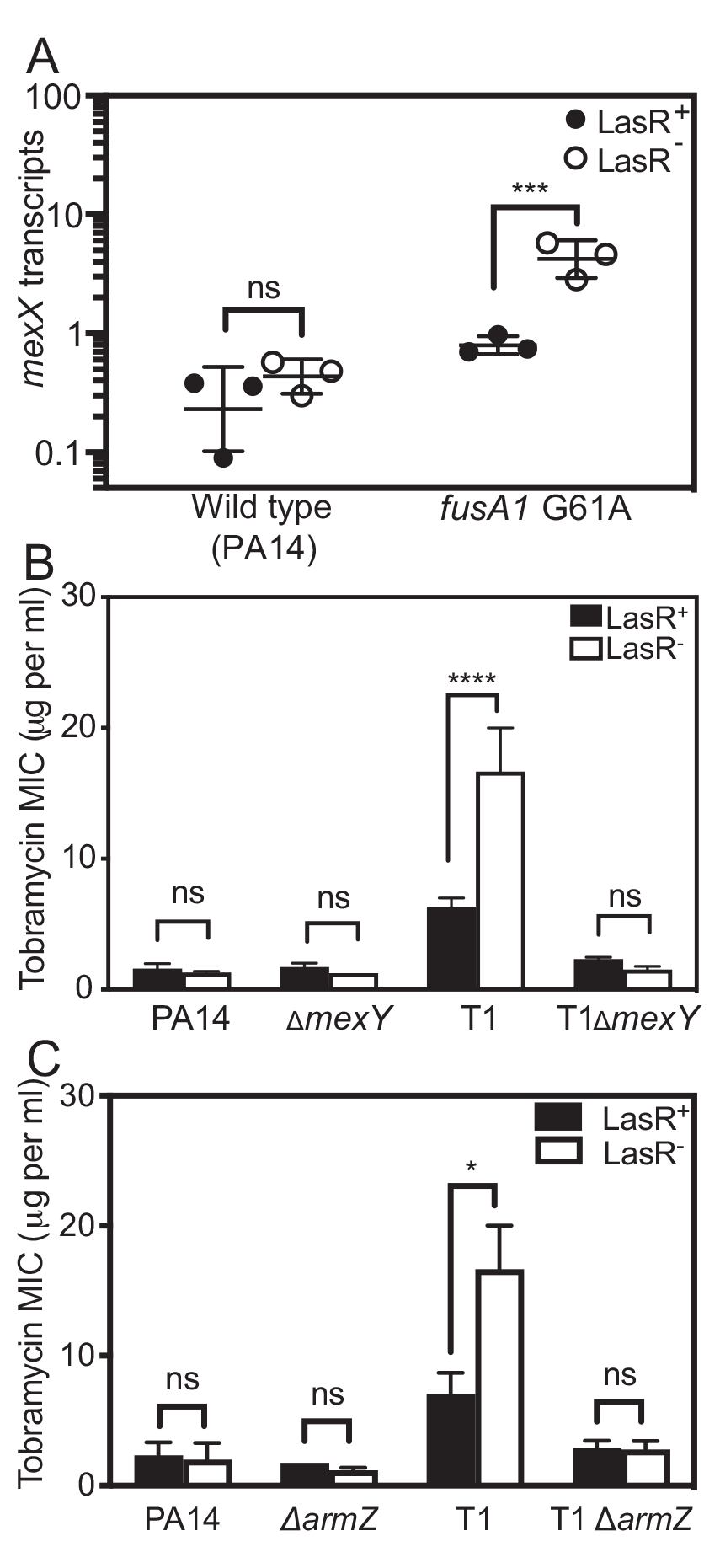
Role of MexXY on tobramycin resistance of Δ*lasR* and *fusA1* G61A mutants. **A.** *mexX* transcript levels were determined by droplet digital PCR and normalized to the housekeeping control gene *proC.* Strains were wild type PA14 or PA14 with the *fusA1* G61A substitution. Each strain had either an intact *lasR* (LasR^+^, filled circles) or Δ*lasR* (LasR^-^, open circles). Each point represents in independent experiment; horizontal lines represent the geometric mean and the vertical lines represent the geometric standard deviation. Statistical analysis by 2-way ANOVA showed a significant interaction between the effects of strain and *lasR* allele on *mexX* transcripts (p<0.005, F_(1, 8)_ = 15.67). **B and C.** The minimum inhibitory concentration (MIC) of tobramycin was determined for each strain carrying intact *lasR* (LasR^+^, black bars) or Δ*lasR* (LasR^-^, white bars). Results are the average of three independent experiments and the vertical bars show standard deviation. In pair-wise comparisons with the LasR^+/-^ PA14 strains there was a significant interaction between the effects of strain and *lasR* allele for T1 (p<0.01 and F_(1,8)_ = 29.01 for panel B and p<0.005 and F(1,8) = 17.99 for panel C) but not for the other strains. In all three panels, pairwise comparisons of LasR^+^ and LasR^-^ of each strain was determined using Sidak’s post-hoc test with p-values adjusted for multiple comparisons. *, p<0.05; **, p<0.01, ***, p < 0.001****, p < 0.0001, ns = not significant.

One mechanism of MexXY induction is through the ArmZ regulator (49, 50), which is activated by a mechanism of transcription attenuation in response to perturbations that cause the ribosome to stall (50-52). ArmZ de-represses the MexXY regulator MexZ; thus activation of ArmZ ultimately leads to increased MexXY activity (49, 50, 53). To test the hypothesis that ArmZ is required for MexXY activation in *fusA1* G61A and Δ*lasR* double mutants, we deleted *armZ* in T1 and the T1 Δ*lasR* mutants and determined the MIC of these strains. Similar to the result of deleting *mexY,* we found that deleting *armZ* abolished LasR-dependent changes in MIC observed in T1 (Fig. 5C). Together, the results support that MexXY and the ArmZ regulator are both required for *fusA1* G61A to increase the tobramycin resistance of Δ*lasR* mutants.

The ArmZ regulator is activated by ribosome stalling, and *fusA1* mutations are associated with slower translation rates (41, 48, 54). Thus, we hypothesized that the *fusA1* G61A mutation might cause ribosome stalling effects that are enhanced in combination with Δ*lasR*. To test this hypothesis, we measured the doubling times of our *fusA1* G61A and Δ*lasR* mutant strains during logarithmic growth in minimal media, as growth rates can be an indirect indication of translation rate. We found that *fusA1* G61A alone had no significant effect on growth, however, in combination with Δ*lasR*, we observed slowed growth (Table 2). These results support the idea that MexXY activation in Δ*lasR* and *fusA1* G61A double mutants might be caused by increased ribosome stalling.

**Table 2.** Growth rates in MOPS minimal medium^a^.

| Strain/Isolate | Doubling time (min)<sup>a, b</sup> | strain | *lasR* | interaction |
| --- | --- | --- | --- | --- |
| PA14 | 62.5 (2.0) |  |  |  |
| PA14 $\Delta lasR$ | 61.4 (2.9) | | | |
| *fusAI* G61A | 66.3 (2.9) | p<0.0005 | p<0.01 | p<0.005 |
| *fusAI* G61A $\Delta lasR$ | 78.2 (2.2)<sup>***, c</sup> | F = 49.91 | F = 13.81 | F = 19.73 |
| T1 | 67.7 (4.0) | p<0.0005 | p<0.02 | p<0.005 |
| T1 $\Delta lasR$ | 79.7 (2.5)<sup>***</sup> | F = 47.61 | F = 10.41 | F = 14.81 |
| T3 | 74.9 (3.4) | p<0.0001 | ns | p<0.04 |
| T3 $\Delta lasR$ | 83.0 (3.5)<sup>*</sup> | F = 96.27 | F = 4.084 | F = 6.930 |

<sup>a</sup>The values represent the average doubling time during logarithmic growth determined from three independent experiments, with the standard deviation indicated in parantheses. Standard deviation was zero where not indicated.
<sup>b</sup> The significance of the effects of strain (PA14, *fusAI* G61A, T1 or T3) and *lasR* allele (intact *lasR* or $\Delta lasR$ ) and the interaction of the two on MIC was determined by 2-way ANOVA by performing pairwise comparisons of each *LasR*<sup>+/−</sup> strain pair with that of PA14. The interaction was significant in all cases. For *fusAI* G61A p<0.005 and F = 19.73; for T1 p<0.005 and F = 14.81; for T3 p<0.04 and F = 6.930.

## DISCUSSION

In this study, we demonstrate that *lasR*-null mutations have strain-dependent effects on resistance to certain antibiotics, which are caused by specific mutations in the translation accessory *fusA1* gene (G61A). These results add to the growing body of work supporting that adaptive mutations can cause unexpected effects on quorum-sensing phenotypes. For example, in a prior study we showed that *ptsP* mutations can influence LasR-dependent cooperation and cheating dynamics (17). In addition, inactivation of the efflux pump regulator MexT enables activation of the RhlR-I system as well as the non-AHL Pseudomonas quinolone signal (PQS) in *lasR*-null mutants (55-57). These are all examples of epistatic gene interactions, where the effect of one gene mutation is modified by mutations in other genes (38). Epistatic interactions have consequences for evolutionary outcomes because they can place constraints on adaptation; for example, some mutations will become favorable only after other mutations have accumulated (58). These studies highlight how epistatic gene interactions can contribute to the evolution of quorum sensing during adaptation to changing environments.

Our finding that *fusA1* G61A mutations modulate the effects of Δ*lasR* on tobramycin resistance are consistent with a type of epistasis called sign epistasis. Sign epistasis is a phenomenon where a trait is beneficial in certain genetic backgrounds and harmful in others (58-60). Under tobramycin selection, Δ*lasR* mutations were deleterious in the laboratory strain PA14 but beneficial in *fusA1* G61A mutant strains (Fig. 3). Sign epistasis has particularly dramatic effects on evolution because it can reverse the effects of selection (61, 62). Sign epistasis has been demonstrated in *P. aeruginosa* in the context of rifampicin resistance (63), where combinations of mutations causing rifampicin resistance were shown to have epistatic effects on fitness in the absence of any antibiotic. About half of the interactions studied showed epistatic effects on fitness, and a surprisingly high number of these (84%) were sign epistatic. Sign epistasis has also been demonstrated in the development of cefotaxime resistance in *Escherichia coli* (64) and the evolution of antibiotic resistance in *Mycobacterium tuberculosis* (65). Such studies illustrate that sign epistatic interactions are strikingly common across a variety of gene mutations and bacterial species. Our findings provide a proof-of-principle that sign epistasis can cause dramatic effects on the evolution of quorum sensing by reversing the effects of antibiotic selection of *lasR* mutants. It will be interesting to determine whether mutations other than those characterized in our study can have similar effects.

The *fusA1* G61A mutation had relatively mild effects in a wild type strain, but more severe growth defects in combination with the *lasR* mutation (Table 2). These results suggest that the deleterious effects of the *fusA1* G61A mutation may be compensated by some unknown function of LasR. In the case of other *fusA1* mutations, it is thought that the mutations in the encoded EF-G1A protein cause the ribosome to remain “stuck” on the mRNA, which might cause stalling when other ribosomes get backed up behind it (66). We posit that LasR activates some function that somehow protects from ribosome stalling. Consistent with FusA1^A21T^ having a defect that is exacerbated by loss of LasR, expression of wild-type, fully functional *fusA1* abrogated the effects of *lasR* deletion on tobramycin resistance in a *fusA1* G61A background (Fig. 2B). Future studies are aimed at understanding how LasR might compensate for the deleterious effects of *fusA1* G61A mutations.

We found it interesting that one other *fusA1* mutation (A1366G) had the opposite effect as the G61A mutation. The *fusA1* A1366G mutation decreased tobramycin resistance of Δ*lasR* mutants (Fig. 4B), whereas the *fusA1* G61A mutation increased resistance. From these results, we posit that LasR may enhance the deleterious effects of the A1366G *fusA1* mutation rather than providing protection as we propose for G61A *fusA1*. Most other *fusA1* mutations did not have interactive effects with *lasR.* These results show that LasR and *fusA1* gene interactions can be neutral, beneficial or deleterious depending on the type of *fusA1* mutation. Our results support the idea that different *fusA1* mutations increase antibiotic resistance through different pathways. Our result that some mutations increase antibiotic sensitivity of Δ*lasR* mutants opens the door to the intriguing possibility of combination therapies that rely on epistatic gene interactions. For example, combining a LasR inhibitor with a therapeutic that enhances antibiotic sensitivity of a *lasR* mutant could be an effective novel approach to therapeutic design.

## MATERIALS AND METHODS

### Culture conditions and reagents

Routine growth was in Luria-Bertani broth (LB) for *Escherichia coli* or in LB buffered to pH 7 with 50 mM 3-(morpholino)-propanesulfonic acid (LB-MOPS) for *Pseudomonas aeruginosa* or on LB agar (LBA; 1.5% wt/vol Bacto Agar). Liquid growth media for specific experiments were M9-caseinate (casein broth); [6 g liter^−1^ Na_2_HPO_4_, 3 g liter^−1^ KH_2_PO_4_, 0.5 g liter^−1^ NaCl, 1 g liter^−1^NH_4_Cl, pH 7.4, 1% sodium caseinate)](17), or a MOPS minimal medium [25 mM d-glucose, freshly prepared 5 μM FeSO_4_, 15 mM NH_4_Cl, and 2 mM K_2_HPO_4_ added to a 1× MOPS base buffer consisting of 50 mM MOPS, 4 mM Tricine, 50 mM NaCl, 1 mM K_2_SO_4_, 50 μM MgCl_2_, 10 μM CaCl_2_, 0.3 μM (NH_4_)_6_Mo_7_O_24_, 40 μM H_3_BO_3_, 3 μM cobalt(II) acetate, 1 μM CuSO_4_, 8 μM MnSO_4_, and 1 μM ZnSO_4_] (17, 67, 68). 4% skim milk agar (SMA) was used for identifying Δ*lasR* mutants in competition experiments. All growth was at 37°C and liquid cultures were grown with shaking at 250 rpm in 18 mm culture tubes (2-ml cultures), 125 ml baffled flasks (10-ml cultures) or 250-ml baffled flasks (60-ml cultures). For strain construction, we used 100 µg ml^-1^ carbenicillin, 15 µg ml^-1^ gentamicin, and 2.5–10 µg ml^-1^ tetracycline for *E. coli* and 150–300 µg ml^-1^ carbenicillin, 50–200 µg ml^-1^ gentamicin, and 15–100 µg ml^-1^ tetracycline for *P. aeruginosa.* 3OC12-HSL was purchased from Cayman Chemicals (MI, USA), dissolved in acidified ethyl acetate with glacial acetic acid (0.1 ml l^−1^) and added to an empty sterile conical tube and dried by evaporation prior to adding liquid media.

Genomic or plasmid DNA was extracted using Qiagen Puregene Core A kit (Hilden, Germany) or IBI Scientific plasmid purification mini-prep kit (IA, USA) while PCR products were purified using IBI Scientific PCR clean-up/gel extraction kits, according to the manufacturer’s protocol. Antibiotics were purchased from GoldBio (MO, USA) except for tetracycline, which is from Fisher Scientific (PA, USA).

### Bacterial strains and strain construction

All bacterial strains, plasmids, and primers used in this study are provided in the Supplementary Material. *P*. *aeruginosa* strain UCBPP-PA14 (‘PA14’) and PA14 derivatives were used for these studies. Markerless deletions in specific loci of *P*. *aeruginosa* PA14 were generated using allelic exchange as described previously (69). To generate plasmids for allelic exchange, DNA fragments with the mutated or deleted gene allele plus 500 bp flanking DNA were synthesized (Genscript, New Jersey USA) or generated by PCR using primer-incorporated restriction enzyme sites. The synthesized or PCR product were moved to plasmid pEXG2 (70) using restriction enzyme digestion and ligation or isothermal assembly. The subsequent plasmids were transformed into the appropriate *P. aeruginosa* strain using described methods (71). The plasmids for Δ*lasR* (68), Δ*mexY* (72), and Δ*armZ* (53) were described elsewhere. Merodiploids were selected on Pseudomonas Isolation Agar (PIA)-carbenicillin (150–300 μg ml^-1^) for Δ*lasR*; PIA-gentamicin (50–200 μg ml^-1^) for *fusA1* G61A, Δ*lasI*, and *ptsP* 1547T; and PIA-tetracycline (15–100 µg ml^-1^) for Δ*mexY* and Δ*armZ*. Deletion mutants were counterselected using NaCl-free 15% sucrose. Putative mutants were verified through antibiotic sensitivity tests and gene-targeted Sanger sequencing. To make the P*rha*-*fusA1* expression cassette, we PCR-amplified the wild-type *fusA1* gene from PA14 using primer-encoded restriction enzyme sites. The PCR product was digested and ligated to pJM253 (miniCTX1-rhaSR-PrhaBAD)(73). This plasmid was moved into *P. aeruginosa* by conjugation as described previously (73). Transformants were selected on LBA with 200 µg ml^-1^ tetracycline and the insertion of the cassette in the *attB* site was verified by PCR.

### Antimicrobial susceptibility assays

Antibiotic susceptibility was determined by MIC using a modified method from the 2022 guidelines of the Clinical and Laboratory Standards Institute (CSLI) (74), similar to that previously described (17). Briefly, two antibiotic dilution series were made from staggered starting antibiotic concentrations to cover a broader range of concentrations in MOPS minimal medium, and successively diluted 2-fold in a 200 µl volume in 2 ml tubes. The starter cultures were prepared by growing *P. aeruginosa* in LB-MOPS to an optical density at 600 nm (OD_600_) of 4. The starter cultures were subsequently diluted 1:40 into each tube containing tobramycin to start the MIC experiment. After 20 h of incubation with shaking, turbidity was measured using a Biotek Synergy 2 plate reader. The MIC was defined as the lowest concentration of tobramycin (µg ml^-1^) in which bacterial growth was not measurable.

### Coculture assays

Overnight (18 h) pure cultures were grown in LB-MOPS, diluted to an OD_600_ of 0.025 for LasR^-^ or 0.05–0.15 for LasR^+^ into LB-MOPS, and grown to an OD_600_ of ∼3.5 before combining at a 99:1 (LasR^+^: LasR^-^) ratio and used to start the coculture by diluting 1:40 into casein broth in 18 mm test tubes. In some cases as indicated, tobramycin was added to the casein broth coculture. At 24 h, cocultures were diluted 1:40 into fresh casein broth in a new test tube and the experiment was ended at 48 h. The initial and final total population counts (CFU ml^-1^) were determined by dilution plating and viable plate counts. The % Δ*lasR* mutant was determined by patching 200 colonies on SMA where Δ*lasR* mutants form distinct colony phenotypes (17, 22, 23, 75).

### Droplet digital PCR

Overnight (18 h) pure cultures were grown in LB-MOPS, subcultured in LB-MOPS, and grown to an OD_600_ of ∼4. Stationary-phase *P. aeruginosa* (OD_600_ of 4) was diluted to OD_600_ 0.1 in MOPS minimal medium and grown 2.5-3 hours (OD_600_ ∼0.20-0.45). RNA was harvested using the RNeasy minikit (Qiagen), following the manufacturer’s instructions. Droplet digital PCR (ddPCR) was performed on a Bio-Rad QX200 ddPCR system using Eva Green supermix. Each reaction mixture contained 1.8 ng µl^-1^ cDNA template, 0.8 µM of each primer, and 10 µl Eva Green supermix in a 20-µl final volume. After generating 40 µl of oil droplets, 40 rounds of PCR were conducted using the following cycling conditions: 95°C for 30s, 62°C for 30s, and 68°C for 30s. Absolute transcript levels were determined using Bio-Rad QuantaSoft software. In all cases, a no-template control was run with no detectable transcripts. The reference gene used was proline biosynthetic gene (*proC*), and the results are reported as the calculated transcript amount of a given gene per calculated *proC* transcript.

### Growth Curve

Overnight (18 h) pure cultures were grown in LB-MOPS, subcultured in LB-MOPS, and grown to an OD_600_ of ∼4. Stationary-phase *P. aeruginosa* (OD_600_ of 4) was diluted to OD_600_ 0.1 in MOPS minimal medium, and OD_600_ was measured in a Jenway spectrophotometer every hour for 8 h. An exponential-fit trendline was fit to the data used to calculate the doubling time.

### Statistical analysis

All statistical analyses were carried out using GraphPad Prism version 9.4.0 (GraphPad Software, San Diego, CA). Unless otherwise noted, antibiotic MICs, cocultures, *mexX* transcripts and growth rates were analyzed using 2-way analysis of variance (ANOVA). Significance of LasR- or antibiotic-dependent effects among strains was determined by finding the interaction term with alpha = 0.05 in pair-wise comparisons of each strain/isolate with PA14. Significance of differences between LasR^+^ and LasR^-^ within each strain was determined using Sidak’s multiple comparisons test in a posthoc analysis. Statistically significant differences are defined in the figure legends.

## ACKNOWLEDGEMENTS

This work was supported by the NIH through grant R35GM133572, CMADP COBRE (P20 GM103638), and K-INBRE (P20 GM103418) and by Inez Jay Fund to J.R.C. V.D.C. was supported by an Undergraduate Research Award from the KU Center for Undergraduate Research and a K-INBRE fellowship (P20 GM103418). K.A.T. was supported by KU Center for Undergraduate research Emerging Scholars program, U.S. Department of Education McNair Scholars Program, and Maximizing Access to Research Careers (MARC) (T34GM136453-01).

R.G.A. was supported by the Fulbright Foreign Student Program (15160174). A.H. was supported by the NIH Bridges to Baccalaureate Program (R25 GM060182). The authors would also like to acknowledge Dr. Keith Poole (Queen’s University) and Dr. Katy Jeannot ((Université de Franche-Comté) for providing plasmids; Nicole E. Smalley and Dr. Ajai A. Dandekar (University of Washington), and Dr. Robert Unckless (University of Kansas) for the insightful suggestions; and Rishita Yadali, Isabelle Parisi, Emma Norris, and Benjamin Smith for their technical support. Molecular graphics and analyses performed with UCSF ChimeraX, developed by the Resource for Biocomputing, Visualization, and Informatics at the University of California, San Francisco, with support from National Institutes of Health R01-GM129325 and the Office of Cyber Infrastructure and Computational Biology, National Institute of Allergy and Infectious Diseases.

