## Supplemental for "Tobramycin adaptation alters the antibiotic susceptibility of *Pseudomonas aeruginosa* quorum sensing-null mutants"

^§^Contributed equally

^b^ Present address: Rhea G. Abisado-Duque, Department of Biology, Ateneo de Manila University, Quezon City, Philippines

**Table S1.** Mutations in isolates from tobramycin-treated populations^a^.

| Isolate | Gene mutation^b^ | | |
| --- | --- | --- | --- |
|  | *ptsP* | *fusA1* | others |
| T1 | 1547T | G61A | *pmrB, mdpA* |
| T2 | 841ΔC | ND^c^ | *mexZ*, *rrf2*, *psdR,* PA14_RS15595 |
| T3  T4  T5  T6 | 1547T  **G392228A^d^**  **G392228A^d^**  **A392231G^d^** | G61A  A1655G  G1634A  ND | *dppA3*, *ostA*  *psdR*  *mdpA*  *mdpA, pstS*, *ftsH*, *nuoM*, Δ**4064707-4092544** |

^a^Table adapted from Abisado et al. (1), in which isolation and genome sequencing are described in detail. In brief, isolates were from populations passaged with tobramycin at 0.6-7.1 µg ml^-1^ (T1-T3) or 0.6 µg ml^-1^ (T4-T6).

^b^Bolding indicates promoter mutations and large chromosomal deletions that are given by genomic location; all other gene mutations are given by nucleotide location.

^c^ND: no gene mutations were detected

^d^Indicates DNA sequence in the predicted promoter of *ygdP,* which is immediately upstream of *ptsP* and predicted to be co-transcribed with *ygdP* (1).

**Table S2.** Strains carrying *fusA1* G61A mutation^a, b^

Strain	RefSeq	Source & location	Ref.	(date isolated)

PAC14B IPC137_22.1	NZ_RWSP01000022.1	Cystic fibrosis patient	(2)

Sherbrook Canada (2008)

5992 IPC1315_14.1	NZ_RWOU01000014.1	Cystic fibrosis patient	(2)

Quebec City Canada (2014)

20 IPC1603_22.1	NZ_RWLA01000022.1	Cystic fibrosis patient	(2)

Saguaney Canada (2016)

FLR01 scaffold55.1	NZ_PXNR01000055	Cystic fibrosis patient	(3)

San Diego USA (2013)

^a^4,312 total unique *P. aeruginosa* strains with *fusA1* were identified from the pseudomonas.com database (4). These were searched for mutations in amino acid 21 of EF-G1A (encoded by *fusA1*) using Unipro UGENE.

^b^All strains listed carry the G61A nucleotide mutation causing A21T amino acid mutation. No other strains carrying the same amino acid mutation were identified.

**Table S3.** Bacterial strains used in this study.

Strain	Relevant properties	Reference or source

*P. aeruginosa strains*

UCBPP-PA14	Ancestral wild type	(5)

PA14 Δ*lasR*	PA14 with a deletion of *lasR*	(6)

PA14 *fusA1* G61A	PA14 with the *fusA1* G61A mutation	This study

PA14 *fusA1* G61A Δ*lasR*	PA14 *fusA1* G61A with a deletion of *lasR*	This study

PA14 *ptsP* 1547T	PA14 with the *ptsP* 1547T mutation	This study

PA14 *ptsP* 1547T Δ*lasR*	PA14 *ptsP* 1547T with a deletion of *lasR*	This study

PA14 Δ*mexY*	PA14 with a deletion of *mexY*	This study

PA14 Δ*lasR* Δ*mexY*	PA14 Δ*lasR* with a deletion of *mexY*	This study

PA14 Δ*armZ*	PA14 with a deletion of *armZ*	This study

PA14 Δ*lasR* Δ*armZ*	PA14 Δ*lasR* with a deletion of *armZ*	This study

PA14 *attB*::P*rha*	Made using pJM253 (pCTX-1 P*rha*)	This study

PA14 Δ*lasR attB*::*Prha*	Made using pJM253	This study

PA14 *fusA1* G61A *attB*::*Prha*	Made using pJM253	This study

PA14 *fusA1* G61A Δ*lasR attB*::*Prha*	Made using pJM253	This study

PA14 *fusA1* G61A *attB*::*Prha*-*fusA1* Made using pCTX-1 P*rha*-*fusA1*	This study

PA14 *fusA1* G61A Δ*lasR* *attB*::*Prha*-*fusA1*	Made using pCTX-1 P*rha*-*fusA1*	This study

PA14 *fusA1* T119A	PA14 with the *fusA1* T119A mutation	This study

PA14 *fusA1* G352A	PA14 with the *fusA1* G352A mutation	This study

PA14 *fusA1* A1366G	PA14 with the *fusA1* A1366G mutation	This study

PA14 *fusA1* C1664A	PA14 with the *fusA1* C1664A mutation	This study

PA14 *fusA1* A2011G	PA14 with the *fusA1* A2011G mutation	This study

PA14 *fusA1* T119A Δ*lasR*	PA14 *fusA1* T119A with a deletion of *lasR*	This study

PA14 *fusA1* G352A Δ*lasR*	PA14 *fusA1* G352A with a deletion of *lasR*	This study

PA14 *fusA1* A1366G Δ*lasR*	PA14 *fusA1* A1366G with a deletion of *lasR*	This study

PA14 *fusA1* C1664A Δ*lasR*	PA14 *fusA1* C1664A with a deletion of *lasR*	This study

PA14 *fusA1* A2011G Δ*lasR*	PA14 *fusA1* A2011G with a deletion of *lasR*	This study

*PA14 isolates from tobramycin passaging experiment*

T1	Isolate from experiment with tobramycin added at 0.6–7.1µg ml^-1^	(6)

T2	Isolate from experiment with tobramycin added at 0.6–7.1µg ml^-1^	(6)

T3	Isolate from experiment with tobramycin added at 0.6–7.1µg ml^-1^	(6)

T4	Isolate from experiment with tobramycin added at 0.6 µg ml^-1^	(6)

T5	Isolate from experiment with tobramycin added at 0.6 µg ml^-1^	(6)

T6	Isolate from experiment with tobramycin added at 0.6 µg ml^-1^	(6)

*Modified PA14 isolates*

T1 Δ*lasR*	T1 with a deletion of *lasR*	This study

T1 Δ*lasI*	T1 with a deletion of *lasI*	This study

T1 Δ*mexY*	T1 with a deletion of *mexY*	This study

T1 Δ*lasR* Δ*mexY*	T1 Δ*lasR* with a deletion of *mexY*	This study

T1 Δ*armZ*	T1 with a deletion of *armZ*	This study

T1 Δ*lasR* Δ*armZ*	T1 Δ*lasR* with a deletion of *armZ*	This study

T1 *attB*::*Prha*	Made using pCTX-1 P*rha*	This study

T1 *attB*::*Prha*-*fusA1*	Made using pCTX-1 P*rha*-*fusA1*	This study

T2 Δ*lasR*	T2 with a deletion of *lasR*	This study

T3 Δ*lasR*	T3 with a deletion of *lasR*	This study

T4 Δ*lasR*	T4 with a deletion of *lasR*	This study

T5 Δ*lasR*	T5 with a deletion of *lasR*	This study

T6 Δ*lasR*	T6 with a deletion of *lasR*	This study

*Escherichia coli strains*

DH5α	F- Φ80*lacZ* Δ*M15* Δ(*lacZYA*-*argF*) *U169 hsdR17*(rK- mK^+^)	Invitrogen

*recA1 endA1 phoA supE44 thi-1 gyrA96 relA1* λ**^-^**

S17-1	*recA pro hsdR RP4-2-Tc::Mu-km::Tn7*	(7)

SM10	*thi thr leu tonA lacY supE recA*::RP4-2-Tc::Mu Km λ*pi*r	(7)

Rho3	*thi-1 thr-1 leuB26 tonA21 lacY1 supE44 recA*,	(8)

integrated RP4-2 Tcr::Mu (λ*pir* ^+^)Δ*asd*::FRT Δ*aphA*::FRT

HB101	*supE44 hsdS20(r_B_- m_B_-) recA13 ara-14 proA2 lacY1 galK2*	(9)

*rpsL20 xyl-5 mtl-1 leuB6 thi-1*

CC118 λ*pir	Δ(ara-leu) ara D ΔlacX74 galE galK phoA20 thi-1 rpsE rpoB*	(10)

*argE(Am) recAl lysogenized with* λ*pir phage*

**Table S4.** Plasmids used in this study.

Plasmid	Relevant properties	Reference or source

pDA8	pEX18Ap containing PA14 Δ*lasR* with flanking sequences, Ap^R^	(6)

pEXG2	Suicide vector, Gm^r^	(11)

pEXG2-Δ*lasI*	pEXG2 containing PA14 Δ*lasI* with flanking sequences	This study

pEXG2-*fusA1* G61A	pEXG2 containing G61A mutation in *fusA1*	This study

pEXG2-*ptsP* 1547T	pEXG2 containing 1547T mutation in *ptsP*	This study

pCSV05	pEX18Tc containing Δ*mexY* with flanking sequences, Tc^R^	(12)

pYM008	pEX18Tc containing Δ*armZ* with flanking sequences, Tc^R^	(13)

pJM253 (pCTX-1 P*rha*) miniCTX1-rhaSR-PrhaBAD, Tc^R^	(14)

pCTX-1 P*rha*-*fusA1*	pJM253 with the PA14 wild-type *fusA1* allele	This study

pKNG101-*fusA1*_T119A_	pKNG101 containing *fusA1* T119A, Str^R^	(15)

pKNG101-*fusA1*_G352A_	pKNG101 containing *fusA1* G352A, Str^R^	(15)

pKNG101-*fusA1*A_1366G_	pKNG101 containing *fusA1* A1366G, Str^R^	(15)

pKNG101-*fusA1*_C1664A_	pKNG101 containing *fusA1* C1664A, Str^R^	(15)

pKNG101-*fusA1*_A2011G_	pKNG101 containing *fusA1* A2011G, Str^R^	(15)

pRK2013	Helper plasmid for mobilization of non-self-transmissible plasmids; (16)

ColE1 Tra^+^ Mob^+^ Kan^R^

**Table S5.** Primers used in this study.

Primer	Sequence^a^

rgaoligo72^b^	TAATAA**AAGCTT**GTGCTGGCTGATCCGAAATA

rgaoligo73^b^	TAATAA**TCTAGA**GACCGACGAAGTTCTCTTCAG

*Droplet Digital PCR*

mexXF	CGAAGAAGCAGCGGACAC

mexXR	GCAGCTCGCTGGTGATG

proCF	GCCAACGCGCAGGTCAG

proCR	CGGTCACCGCGTCGATCTG

^a^Bolded text indicates restriction sequences

^b^rgaoligo72 and 73 were used to amplify the *fusA1* G61A mutation to construct a vector for allelic replacement in PA14

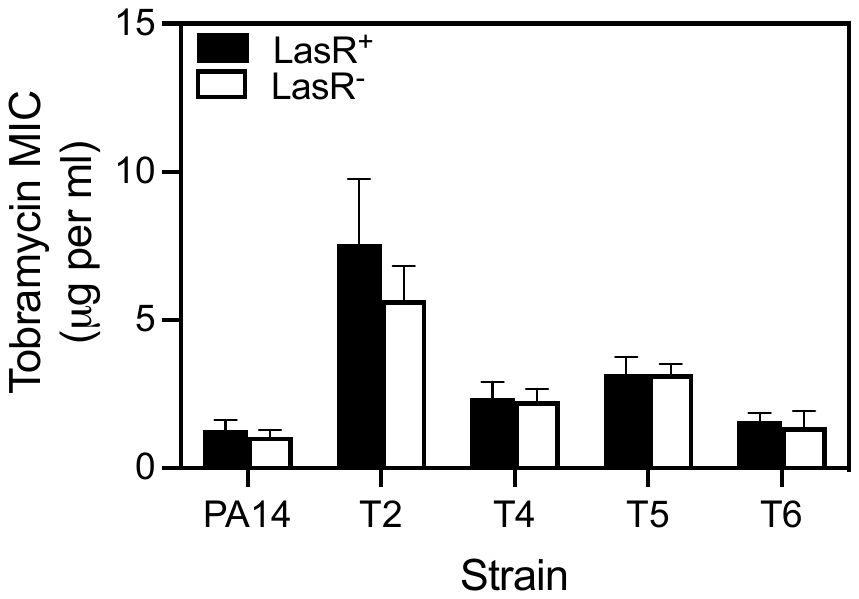

**Fig. S1. Effects of tobramycin on** Δ***lasR* mutants in T2, T4, T5 and T6.** The minimum inhibitory concentration (MIC) of tobramycin was determined for strains carrying the Δ*lasR* mutation (LasR^-^, white bars) or with *lasR* intact (LasR^+^, black bars). Statistical analysis by 2-way ANOVA showed no significance of interaction of *lasR* vs. strain in pairwise comparisons of each isolate with PA14 (p>0.05). There were also no significant differences using Sidak’s post-hoc to directly compare LasR^+^ and LasR^-^ of each strain with p-values adjusted for multiple comparisons of all strains.

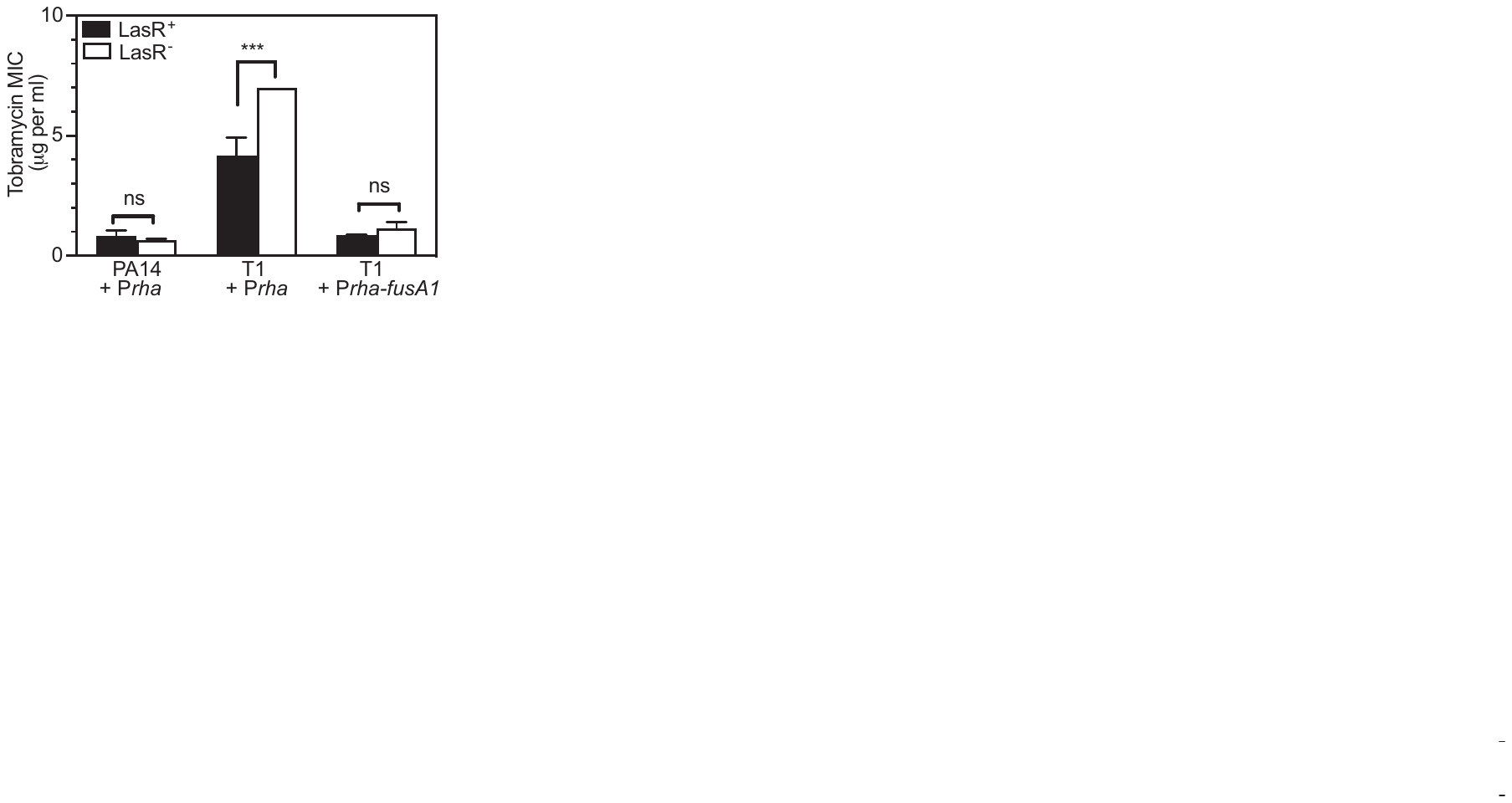

**Fig. S2. Inducible expression of *fusA1* in T1 mutant restores Δ*lasR-*dependent effects on tobramycin resistance.** The minimum inhibitory concentration (MIC) of tobramycin was determined for each strain carrying intact *lasR* (LasR^+^, black bars) or Δ*lasR* (LasR^-^, white bars). Statistical analysis by 2-way ANOVA was used to determine the effect of strain and *lasR* allele and the interaction of the two on MIC of each strain in a pair-wise comparison with PA14. The interaction was significant for T1 + P*rha* (F_1,8_ = 42.76 and p<0.0005); for T1 + P*rha-fusA1* the interaction was not significant (F_1,8_ = 4.884 and p = 0.0581). Comparisons of LasR^+^ and LasR^-^ of each strain was performed using a Sidak’s post-hoc analysis of a 2-way ANOVA of all strains with p-values adjusted for multiple comparisons. For statistical analyses, ***, p<0.001; ns = not significant.
